# Heterogeneous distribution of kinesin-streptavidin complexes revealed by Mass Photometry

**DOI:** 10.1101/2023.12.21.572878

**Authors:** Jing Xu, Nathaniel J. S. Brown, Yeonee Seol, Keir C. Neuman

## Abstract

Kinesin-streptavidin complexes are widely used in microtubule-based active-matter studies. The stoichiometry of the complexes is empirically tuned but experimentally challenging to determine. Here, mass photometry measurements reveal heterogenous distributions of kinesin-streptavidin complexes. Our binding model indicates that heterogeneity arises from both the kinesin-streptavidin mixing ratio and the kinesin-biotinylation efficiency.

---

Kinesin-streptavidin complexes are widely used to drive filament-filament sliding in microtubule-based active matter studies^1-8^. The complexes are typically formed via incubating biotinylated kinesin dimers with streptavidin tetramers. Each kinesin dimer contains up to two biotins^9-11^, and each streptavidin tetramer can bind up to four biotins. The resulting complexes are generally assumed to contain a homogeneous population of two kinesin dimers and one streptavidin tetramer, corresponding to a 2:1 complex stoichiometry. This assumption, however, has not been experimentally verified. Instead, an early study employing analytical gel filtration suggested up to 8 kinesin dimers per complex^2^, and recent work employing dynamic light scattering estimated a stoichiometry of four kinesin dimers per streptavidin tetramer^8^. Furthermore, the assumption of a homogeneous population of 2:1 complex stoichiometry is challenged by experimental findings that the macroscopic dynamics of the active matter system depend sensitively on the mixing ratio of kinesin dimers to streptavidin tetramers^1, 4^. Specifically, under otherwise identical conditions, Henkin et al. reported a ∼2.6-fold linear decrease in the characteristic length scale of the active matter system as the mixing ratio increased by ∼3.8-fold from ∼0.9 to 3.4 kinesin dimers per streptavidin tetramer^4^. Because an increase in the number of kinesins in a complex can increase the number of microtubules dynamically crosslinked together, we hypothesized that the decrease in the characteristic length scale in the active matter system could reflect an increase in the relative abundance of kinesin-streptavidin complexes with stoichiometries exceeding 2:1.

Quantitative characterization of kinesin-streptavidin complex stoichiometry is experimentally challenging. For example, the relatively high concentrations of proteins required for analytical gel filtration or ultracentrifugation can induce non-specific aggregates that are otherwise not present at the sub-µM concentrations employed in active-matter work. Although dynamic light scattering can detect particle sizes at dilute concentrations, the resulting size distribution is strongly sensitive to assumptions of protein shape and is further complicated by the strong dependence of scattering intensity on the individual masses of scattering particles^12, 13^. Native polyacrylamide gel electrophoresis (native PAGE), another important technique for characterizing protein complexes in the native form, is sensitive to the shape and net charge, as well as the molecular mass, of the protein^14^. Finally, fluorescence-based methods, such as stepwise photobleaching, are complicated by incomplete fluorescence-labelling of proteins^15^.

Mass photometry is a recently-developed, label-free, technique for determining the mass and the relative abundance of proteins and protein complexes in dilute solutions^16^. Similar to dynamic light scattering, mass photometry is a light scattering-based technique^17-20^. Distinct from dynamic light scattering that employs the interference between light scattered from distinct particles in solution, mass photometry utilizes the interference between light reflected by the sample surface and light scattered by individual particles on the same surface (Fig. 1a). As the result, for each particle that binds the sample surface from solution, mass photometry returns a concentric ring pattern characteristic of the interference between a plane wave (reflected by the fixed sample surface) and a spherical wave (scattered by the bound particle) (Fig. 1b). The intensity of the plane wave (reflected by the fixed sample surface) is kept constant, yielding a peak interference intensity that is proportional to the polarizability of the bound particle. For simple dielectric materials, the polarizability of a particle scales linearly with the volume and thus the mass of the particle. This linearity between interference intensity and the mass of the particle is demonstrated in mass photometry over a broad mass range of ∼50-5000 kDa^21^. Heterogeneity in interference intensities reveal multiple mass species in solution (Fig. 1b, bottom panel), and molecular counting of different mass species yields the relative abundance of individual mass species in solution.

**Figure 1.**
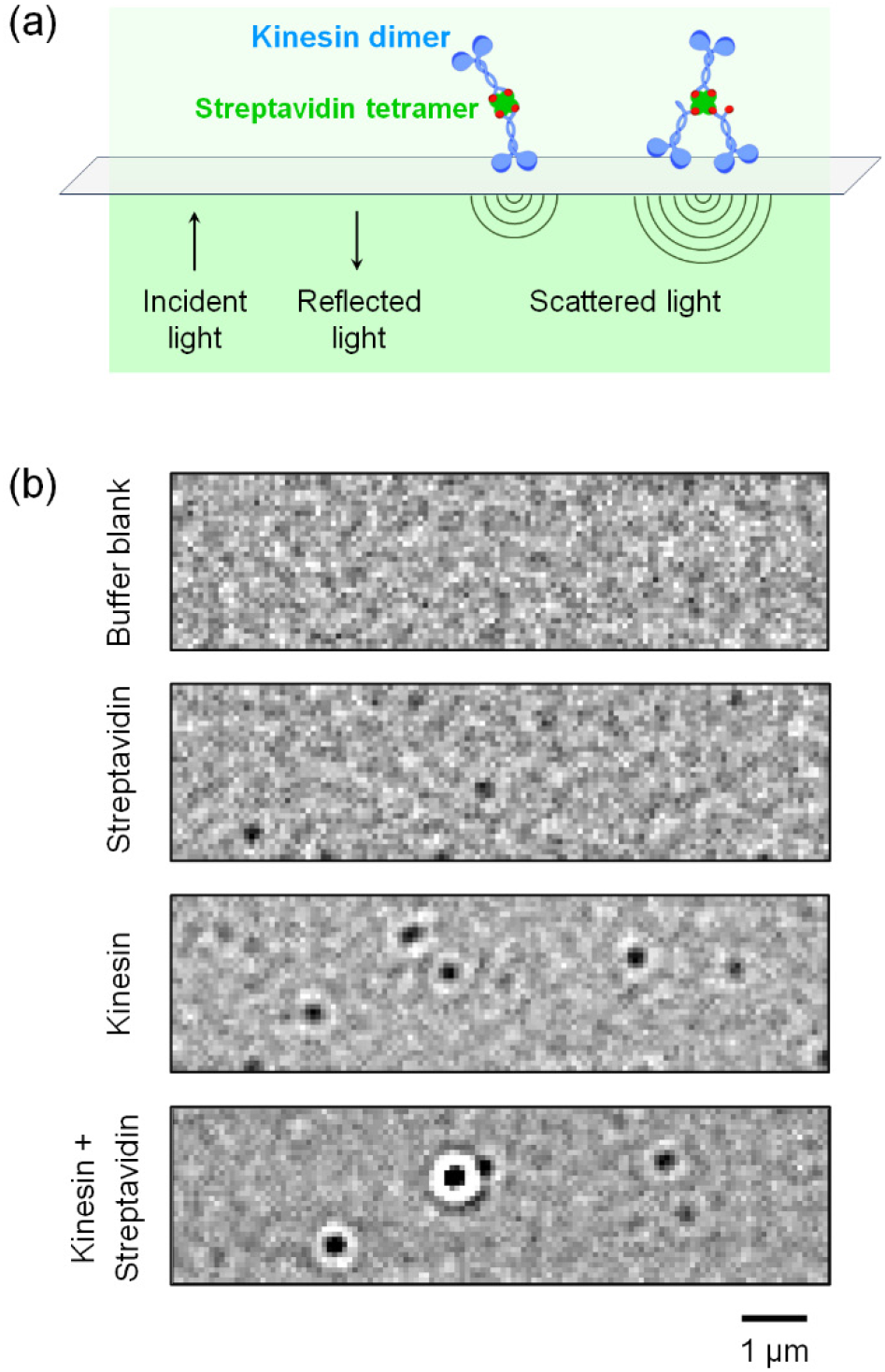
(a) Illustration of mass photometry experiments, in which scattered light from protein complexes on the sample surface interferes with reflected light from the same surface. Illustration is not to scale. (b) Representative mass photometry images of protein-free buffer (top) and protein solutions (bottom three panels). Scale bar, 1 µm. Interference intensity of each spot scales linearly with molecular mass; heterogeneity in spot intensities reveal multiple species of protein complexes in solution (bottom).

In the current study, we carried out mass photometry measurements using a commercial OneMP instrument (Refeyn, UK). We verified that the interference intensity reported by the OneMP instrument scales linearly for masses up to 1048 kDa (Fig. S1, ESI†); this broad linear range is appropriate for determining the molecular mass and thus the stoichiometry of kinesin-streptavidin complexes in the current study. We employed the resulting mass versus interference intensity calibration to convert individual interference intensities from mass photometry images (for example, Fig. 1b) into individual molecular masses.

We prepared kinesin-streptavidin complexes by incubating solution mixtures of kinesin dimers and streptavidin tetramers on ice for 30 min following standard protocols^1, 6, 7^. We varied the kinesin-streptavidin mixing ratio between 0.4-3.6 kinesin dimers per streptavidin tetramer, encompassing the range of mixing ratios previously identified to impact the characteristic length scale in active matter^4^. Note that this prior work^4^ specified the mixing ratios but not the associated protein concentrations. Based on the common range of kinesin concentrations in literature^1, 6, 7^, and to reduce the likelihood of artifactual protein aggregation that can occur at high kinesin concentrations^7^, we limited the concentration of kinesin dimers to ≲2-3 µM for the majority of the measurements in the current study. Accordingly, we employed a constant concentration of streptavidin tetramers (0.6 µM) and varied the concentration of kinesin dimers to achieve the indicated mixing ratio. Given the relatively fast binding rate of biotin to streptavidin (∼3-75 µM^-1^s^−1^)^22, 23^ and the exceedingly slow dissociation rate (∼10^−6^ s^-1^)^24, 25^, we expect kinesin-streptavidin binding to be complete and stable within the 30 min incubation and subsequent measurement time. We estimated the concentrations of the isolated proteins via absorption measurements at 280 nm, and we determined the mixing ratio for each preparation of kinesin-streptavidin complexes via quantitative densitometry of proteins stained with Coomassie blue (Fig. S2, ESI†).

We diluted each preparation of kinesin-streptavidin complexes to 20 nM total protein concentration, within the mass-photometry working concentration range (0.1-100 nM^21, 26^). For each fresh dilution of protein solutions, we imaged the individual binding events for a fixed measurement duration of 1 min. This measurement duration, typical in mass photometry experiments, is short compared to the exceedingly slow dissociation rate of biotin-streptavidin binding (∼10^−6^ s^-1^)^24, 25^. Together, the short measurement duration and the dilute protein concentration preserve complex integrity while avoiding protein aggregation. Solutions containing isolated proteins were used as controls.

We performed 12-22 independent mass photometry measurements for each preparation of kinesin-streptavidin complexes or isolated protein controls. We detected no substantial variations among independent measurements using the same sample (Fig. S3a, ESI†) or using different preparations of the same kinesin-streptavidin mixture (Fig. S3b, ESI†). We pooled these independent measurements to determine the distribution of mass species in each protein solution (Fig. 2). We fitted the resulting mass distributions to a bi-Gaussian mixture model (red lines, Fig. 2) to first determine the molecular masses of the major species (dashed lines i-v, Fig. 2). To ensure fitting accuracy, only mass species with pronounced peak profiles (>40 in peak height, or >400 counts total) were included in the fit (red lines, Fig. 2).

**Figure 2.**
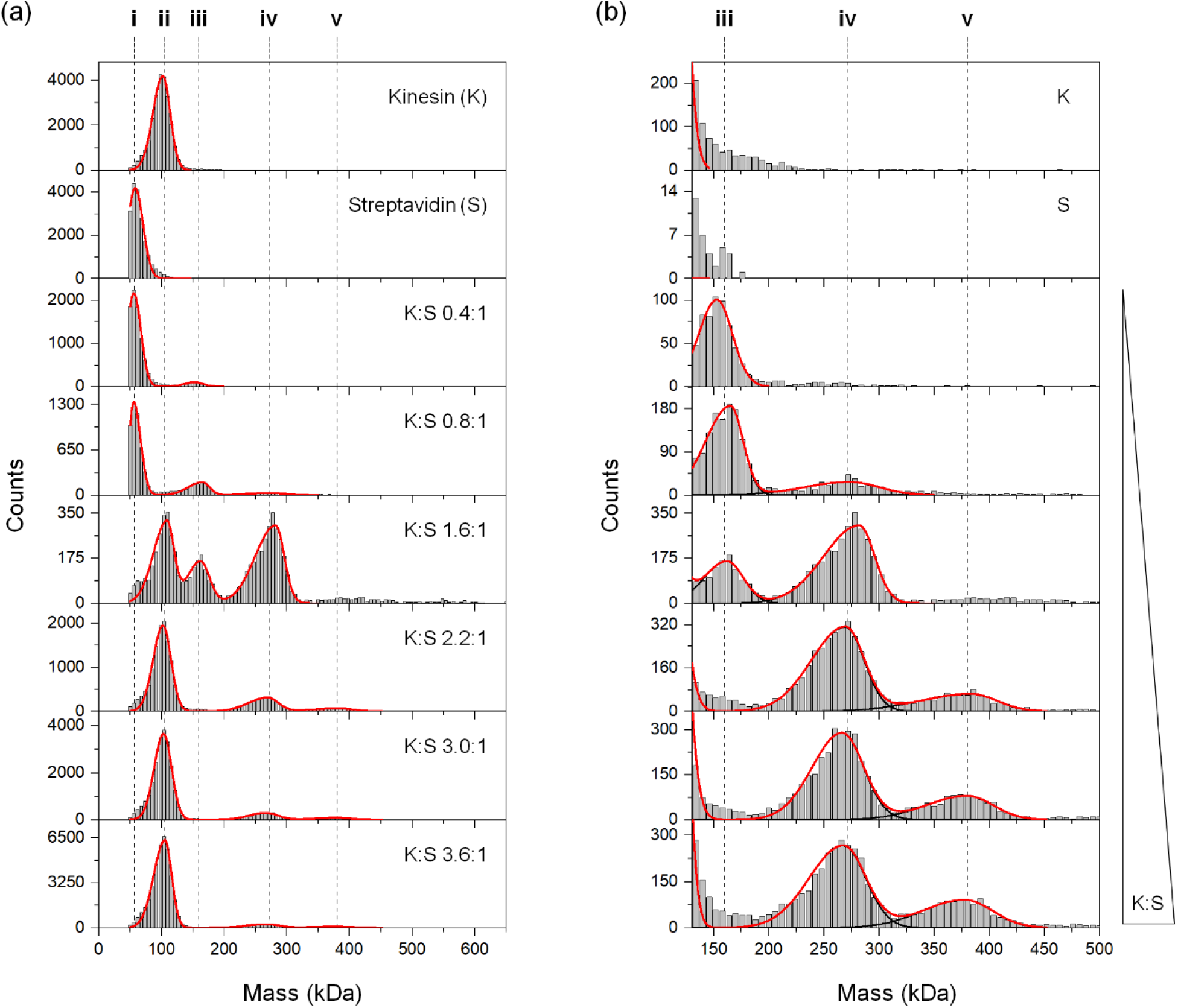
(a) Distributions of molecular masses revealed by mass photometry. Solutions contain kinesin (K), streptavidin (S), and their mixtures (K:S). K:S indicates the molar ratio of kinesin dimers to streptavidin tetramers in each mixture (“mixing ratio”). The concentration of streptavidin in each mixture was kept constant at 0.6 µM. Each distribution represents data pooled from 12-22 independent measurements. Dashed lines indicate molecular masses of identified major species; red lines indicate best fits of mass distributions to a bi-Gaussian mixture model (Materials and Methods, ESI†). To ensure fitting accuracy, only mass species with pronounced peak profile (>40 in peak height, or >400 counts) were included in the fit. (b) Expanded view of mass species iii-v. Red lines and dashed lines are as described in (a). Black lines indicate contributions of individual species to the fit.

We detected single mass species in control solutions of isolated proteins (top two panels, Fig. 2a). The molecular masses of each species are in good agreement with the theoretical masses of the streptavidin tetramer and the kinesin dimer (i-ii, Fig. 3), as well as a prior mass photometry measurement of the streptavidin tetramer (57.2 ± 1.5 kDa in the current study *vs*. 55.7 ± 1.1 kDa previously^16^). These results validate the accuracy of mass photometry and demonstrate no significant protein aggregates in either protein solution.

**Figure 3.**
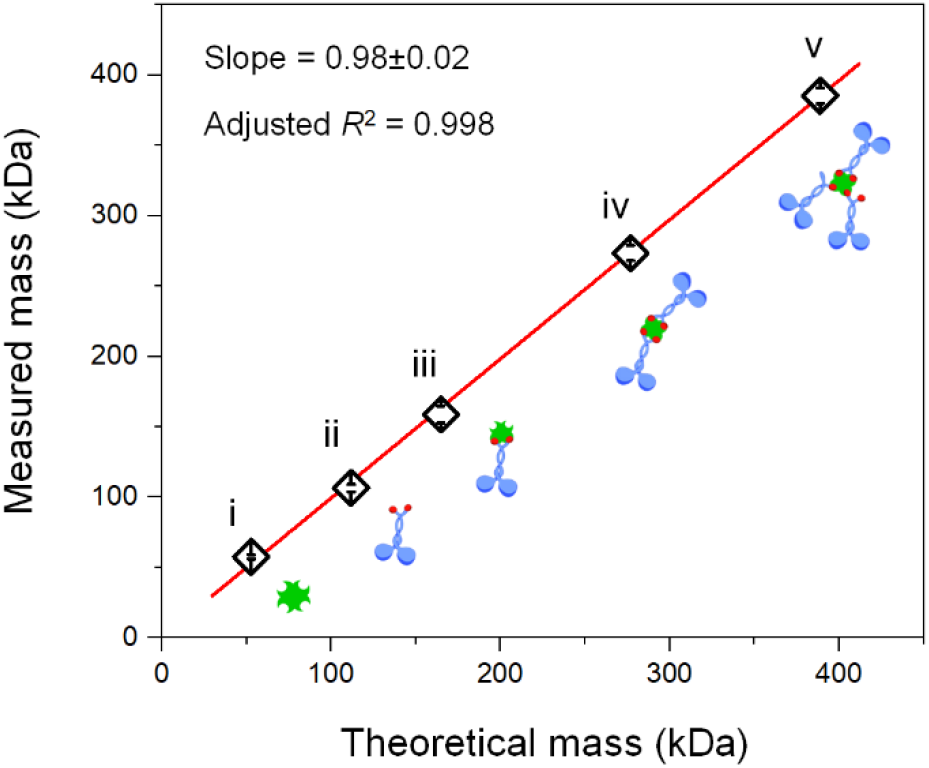
Theoretical *vs*. measured masses for major species revealed by mass photometry. Red line indicates best linear fit with zero intercept. slope = 0.98±0.02 and adjusted *R*^2^ = 0.998. Cartoons illustrate streptavidin (i), kinesin (ii), and kinesin-streptavidin complexes with 1:1, 2:1, and 3:1 stoichiometry (iii, iv, and v, respectively). Measured masses were determined via the best-fits in Fig. 2 (dash lines i-v). Error bars indicate standard deviation. *N* = 4-12.

In contrast, we detected multiple mass species in solutions containing both kinesin and streptavidin (bottom six panels, Figs. 2a and 2b). In addition to the two isolated proteins, mass photometry revealed three larger species (iii-v, Figs. 2a and 2b). The molecular masses of these larger species are highly correlated with theoretical molecular masses for complexes containing up to three kinesin dimers and one streptavidin tetramer (iii-v, Fig. 3). Note that the correspondence between mass and complex stoichiometry is not necessarily unique, given that the molecular mass of the kinesin dimer is approximately twice that of a streptavidin tetramer. For example, the measured mass of species iv could, in principle, be consistent with a complex containing one kinesin dimer and three streptavidin tetramers (∼271 kDa). However, this possibility is unlikely given that the complex was observed in solutions with excess kinesin (Fig. 2b). In addition to complex species iii-v, we also detected substantially larger species that we refer to as “higher-order complexes” (Fig. S4, ESI†). These higher-order complexes likely comprise a mixture of different species: the mass at ∼500 kDa is consistent with a report of four kinesin dimers per streptavidin tetramer^8^, and the mass at ∼950 kDa is consistent with a report of up to 8 kinesin dimers per complex^2^.

We found that the isolated kinesin protein is largely absent in mixtures at lower molar mixing ratios (for example, K:S 0.4:1, Fig. 2a). This finding indicates that all kinesin bound streptavidin during the 30 min incubation. Similarly, the isolated streptavidin protein is absent in mixtures at higher mixing ratios (for example, K:S 3.6:1, Fig. 2a), indicating that all streptavidin bound kinesin.

For most incubation ratios tested, we detected mainly isolated proteins rather than kinesin-streptavidin complexes (Fig. 2a). This observation reflects an important caveat in using mass photometry for molecular counting and abundance measurements. Briefly, mass photometry employs a differential detection method that is sensitive to proteins binding to, and unbinding from, the sample surface^16-20^. Accurate molecular counting is predicated on proteins irreversibly bound to the surface, such that the binding is counted once and only once. If the protein unbinds, the unbinding leads to a change in interference intensity that is equal in magnitude but opposite in sign as a binding event, resulting in an apparent negative mass reading. Moreover, the unbound protein can be counted multiple times, through repeated rebinding and unbinding events, artifactually increasing the molecular counts^21^. Because we detected substantial negative mass counts for the isolated proteins (i-ii, Fig. S5, ESI†), but not kinesin-streptavidin complexes (iii-v, Fig. S5, ESI†), we excluded the isolated proteins from calculations of relative abundance of complex species in the next section.

Focusing on the region of complex species (Fig. 2b), we observed more than one complex species in most kinesin-streptavidin solutions (bottom five panels, Fig. 2b). We determined the abundance of each complex species via best-fits to a bi-Gaussian mixture model (for example, Fig. S6a, ESI†). For complex species without a pronounced peak profile, we employed our measurements of molecular masses in Fig. 3 to constrain their peak positions in the fit. Additionally, for higher order complexes (Fig. S4, ESI†), we estimated their abundance as the cumulative counts of masses ≥450 kDa. Based on these abundance calculations, and excluding the isolated proteins, we determined the relative abundance of different kinesin-streptavidin complexes as a function of mixing ratio (Fig. 4). The resulting relative-abundance calculations agreed well between experiments using different preparations of the same kinesin-streptavidin mixture (Fig. S6, ESI†).

**Figure 4.**
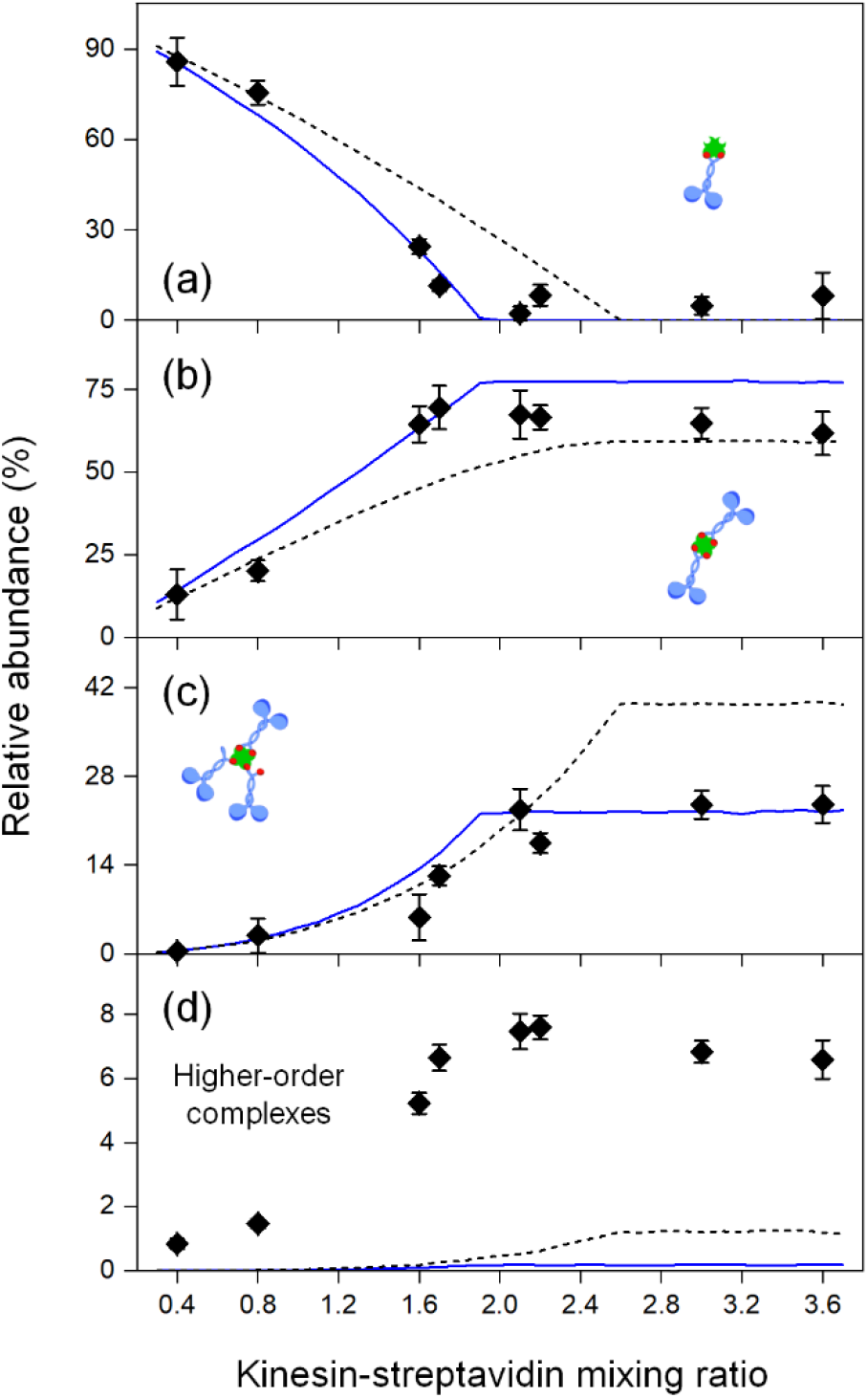
Relative abundance of different complex species as a function of the kinesin-streptavidin mixing ratio. The concentration of streptavidin in each mixture was kept constant at 0.6 µM. Calculations of relative abundance excluded the isolated proteins. (a-c) Complexes with 1:1, 2:1, and 3:1 stoichiometry. Relative abundances were determined via best-fits of mass distributions to a bi-Gaussian mixture model; error bars indicate the associated fitting uncertainties. Black dashed lines indicate the predictions of a simple binding model without any fitting parameters, using the experimentally estimated kinesin-biotinylation efficiency (77%) and kinesin-streptavidin mixing ratios (as indicated). Blue solid lines indicate the predictions of the same binding model, using the best-fitted values of the kinesin-biotinylation efficiency (88%) and an overall scaling factor for the mixing ratios (1.19); only complexes with well-defined stoichiometries (panels a-c) were included in the fit. (d) Higher-order complexes. Relative abundances were estimated as the cumulative counts of masses ≥450 kDa; error bars indicate the associated counting noise. Black dashed lines (and blue solid lines) indicate the binding model predictions using the measured (and the best-fit) values of kinesin-biotinylation efficiency and kinesin-streptavidin mixing ratio as described in (a-c). Note the changes in the y-axis range in panels (a)-(d).

At lower mixing ratios, we found that the most abundant complex consisted of one kinesin dimer and one streptavidin (Fig. 4a), which is functionally equivalent to a single kinesin dimer and cannot drive pair-wise sliding of filaments in active matter. As the mixing ratio increased, the relative abundance of complexes with two or more kinesins increased (Fig. 4b-d). These larger complex species are appropriate for driving active matter dynamics. Moreover, because complexes with three or more kinesin dimers (Fig. 4c-d) can locally crosslink three or more microtubules together, the observed increase in their relative abundance is consistent with a decrease in the characteristic length scale in the active matter, supporting our hypothesis.

We found that changes in the relative abundance of complex species largely plateaued at a mixing ratio of ∼2 kinesin dimers per streptavidin tetramer (Fig. 4). Above this mixing ratio, the relative abundance of complex species varied somewhat, but these variations were within measurement uncertainties (Fig. 4a-c) or did not exceed ∼2% of the overall complex population (Fig. 4d). These observations suggest a more limited range of mixing ratios would impact active-matter length scale than previously found by Henkin et al.^4^ (up to ∼3.4 kinesin dimers per streptavidin tetramer).

Because each streptavidin tetramer contained only four binding sites, plateaus in the relative abundances of complex species is expected. If each kinesin dimer carried two biotin tags (100% biotinylation), we would expect a homogeneous population of complexes with 2:1 stoichiometry at the plateau. In contrast, we observed a heterogenous population of complexes containing two or more kinesins (Fig. 4b-d). Moreover, the substantial presence of three-kinesin complex (Fig. 4c) indicates that some kinesin dimers bound the streptavidin protein via a single biotin.

Employing liquid chromatography-mass spectrometry, we estimated that ∼77% of kinesin monomers in the current study were biotinylated (Fig. S7b, ESI†). Note that, here we employed a standard, biotin carboxyl carrier protein-based approach to attach a single biotin tag on each kinesin monomer^9, 10^; the efficiency of biotinylation using this approach typically ranges between 50 and 80%^11^. To our knowledge, quantitative characterization of kinesin biotinylation has not been reported in prior active-matter studies.

We found that a simple binding model taking account of the kinesin-biotinylation efficiency, and kinesin-streptavidin mixing ratio, captures the main features of the observed complex distributions (Fig. 4). In this binding model, we assumed complete and stable kinesin-streptavidin binding. We assumed that the two biotins on the same kinesin dimer are coupled: once one biotin bound the streptavidin, the second biotin would also bind, provided that there was an open site on the streptavidin. We assumed that a kinesin dimer with two biotins was twice as likely to encounter a streptavidin molecule than a single biotin. We used the molar ratio of biotins and streptavidin monomers in the mixture to estimate the probability that a biotin on a kinesin dimer binds a streptavidin at each encounter. We assumed no steric hinderance between multiple kinesin dimers binding to the same streptavidin tetramer. We considered complexes with well-defined stoichiometries (containing up to four kinesin dimers). We did not consider higher-order complexes.

Using the experimentally estimated kinesin-biotinylation efficiency and kinesin-streptavidin mixing ratios, the model recovered the heterogeneous nature of the complex population, captured the initial changes in the complex heterogeneity as the mixing ratio increased, and closely approximated the relative abundance of two-kinesin complexes in the plateau (black dashed lines, Fig. 4). The predicted presence of three-kinesin complex and higher-order complexes, however, deviated from experiments in their respective plateau values (Fig. 4c-d). These deviations could reflect aggregation into high-order complexes that was not considered in the model. There may also be steric hinderance between multiple kinesins binding to the same streptavidin, which would decrease the relative abundance of the three-kinesin complexes and increase the relative abundance of the smaller complexes. Another deviation from experiments is that the model predicted that the complex heterogeneity plateaued at a mixing ratio ∼30% higher than that estimated in experiments (Fig. 4). This deviation may reflect uncertainties in quantitative determination of the molar mixing ratio in microvolume samples.

To account for measurement uncertainties inherent in the determination of protein mixing ratios and biotinylation efficiency, we varied the efficiency of kinesin biotinylation, and an overall scaling of the mixing ratio, in our model. The resulting best-fit model provided closer approximations to experiments (blue solid lines, Fig. 4), with best-fitted parameters that were within ∼20% of their experimental values (Fig. S8, ESI†). Note that, higher-order complexes were not included in the model or the fit, and the deviation between experiments and model predictions (Fig. 4d) may reflect some aggregation into higher-order complexes.

A simplification in our binding model is that it considers the kinesin-streptavidin mixing ratio, rather than the concentrations of each protein. This simplification is supported by the overall agreement between model predictions and experiments in Fig. 4. We also performed additional experiments in which we kept the mixing ratio constant while we varied the concentrations of kinesin and streptavidin proteins over an 8-fold range (Fig. S9, ESI†). We found that, although the relative abundance of complex species varied somewhat with protein concentration, our model is appropriate for the protein concentrations employed in the current study (Fig. S9, ESI†), which were chosen based on concentrations commonly used in active matter experiments^1, 6, 7^.

Because it is non-trivial to quantitatively control biotinylation efficiency in protein expression, we employed our simple model to explore in more detail how kinesin-biotinylation efficiency impacts the heterogenous distribution of kinesin-streptavidin complexes (Fig. S10, ESI†). In the limit that each kinesin dimer carries two biotin tags (100% biotinylation efficiency), the model recovers the anticipated homogeneous population of complexes with 2:1 stoichiometry when biotin is in excess (dashed lines, Fig. S10, ESI†). Incomplete kinesin biotinylation gives rises to larger complexes with 3:1 or 4:1 stoichiometry (solid lines, Fig. S10c-d, ESI†). These larger complexes have the potential to crosslink multiple microtubules together locally, which could decrease the macroscopic length scale of the resulting microtubule-based active matter. For a given biotinylation efficiency below 100%, our model predicts that the relative abundance of these larger complexes increased as a function of mixing ratio, before plateauing when biotin is in excess (Fig. S10c-d, ESI†). This prediction is consistent with experiments in the current study (Fig. 4), supporting our hypothesis and providing intuition on how the characteristic length scale of the active matter can decrease as the mixing ratio increases (such as that previously reported in Henkin et al.^4^). As the efficiency of kinesin biotinylation decreased, the relative abundance of the larger complexes (3:1 or 4:1 stoichiometry) plateaued at higher values, and the mixing ratio at which the plateau was reached also increased (Fig. S10c-d, ESI†). These predictions further highlight kinesin-streptavidin biotinylation as a key factor in determining the kinesin-streptavidin complex distribution, providing a possible explanation for the impact of mixing ratio on the characteristic length scale observed in active matter experiments^4^.

## Conclusions

Here we employed mass photometry to determine the stoichiometry of Kinesin-streptavidin complexes widely employed in microtubule-based active matter studies. We found that, contrary to the assumption of a 2:1 complex stoichiometry, populations of kinesin-streptavidin complexes are heterogenous (Figs. 2-4), and that the heterogeneity depends on the kinesin-streptavidin mixing ratio (Figs. 2 and 4). We captured the key features of the measured distributions using a simple binding model (Fig. 4). Our model indicates that the efficiency of kinesin-biotinylation is an unexplored determinant of kinesin-streptavidin complex distribution (Fig. S10, ESI†).

It has long been established that that the kinesin: streptavidin mixing ratio can impact the key characteristics of active matter^1, 4^. Here we found that the relative abundance of complexes containing three or more kinesin dimers depended sensitively on the kinesin-streptavidin mixing ratio (Fig. 4). Because these larger complexes can dynamically crosslink three or more microtubules, the observed changes in the relative abundance of these larger complexes provides a mechanistic explanation for the effect of mixing ratio on the characteristic length scale reported previously^4^. Moreover, our model indicates that the effect of mixing ratio on complex heterogeneity is highly sensitive to the efficiency of kinesin biotinylation (Fig. S10, ESI†). Together, our study supports the hypothesis that the macroscopic length scale of the active matter system correlates negatively with the microscopic stoichiometry of the kinesin-streptavidin complexes. Future investigation may help extend our simple model to consider protein concentration, steric hinderance, and higher-order complex formation.

More generally, in addition to active-matter investigation, the biotin-streptavidin chemistry is widely used to induce complex formation in chemical and biological experiments, as well as in biosensor applications^27^. We anticipate that the mass photometry technique used in the current study may be broadly applicable for elucidating the nature of complex formation that underly these experiments and/or biosensing applications.

## Author contributions

J. X. conceived and designed the study; J. X. and N. J. S. B. performed the experiments and analyzed the data; J. X. and K. C. N. developed the binding model; Y. S. generated the recombinant motor protein; J. X. wrote the manuscript. K. C. N. and Y. S. edited it. All the authors reviewed the paper.

## Acknowledgements

This work was supported by the National Institutes of Health (R15 GM120682 to J. X.) and the Intramural Research Program of the National Heart, Lung, and Blood Institute, National Institute of Health (ZIAHL001056 to K. C. N.).

We thank Di Wu, Grzegorz Piszczek, Duck-Yeon Lee, Yasuharu Takagi, and Adam Fineberg for support and comments. We thank the Biophysical Core Facility and the Biochemistry Core Facility for the use of the OneMP instrument and for providing mass spectroscopy measurements. We thank Zvonimir Dogic, Shibani Dalal, the Brandeis Biomaterials Facility, and the National Science Foundation grant no. DMR 2011846 for providing the K401-BIO-6H plasmid. We thank Esme P. Neuman for illustrations of kinesin dimer. We thank the reviewers for insightful comments and suggestions.

## Electronic Supplementary Information

### Materials and Methods

#### Proteins

Dimeric kinesin K401-BIO-H6 (pWC2, Addgene)^1^ carries the first 401 residues of *Drosophila* kinesin heavy chain linked to an 87-residue C-terminus of *E. coli* biotin carboxyl carrier protein followed by a hexa-histidine tag. The expression and purification of k401-BIO-H6 were performed as previously described^1, 2^. In brief, inoculated 2 L of Terrific Broth containing 100 µg/ml Carbenicillin and 25 µg/ml Chloramphenicol was grown overnight at 22°C after induction. Collected cells were lysed by freeze-thaw cycle and high-pressure emulsification (Microfluidics). Following Ni-NTA affinity purification, dimeric and monomeric forms of the protein were separated by size exclusion chromatography using Superdex-200 (Cytiva)^3, 4^. The purity of kinesin proteins from purification was confirmed with denaturing SDS-PAGE shown in Figs. S2a and S7a. The efficiency of biotinylation of kinesin monomer was determined as ∼77% using liquid chromatography-mass spectrometry (Agilent 6224 Accurate-Mass Time-of-Flight LC/MS) (Fig. S7b).

Streptavidin was purchased from Thermo Fisher (catalog # 21122). The purity of streptavidin proteins was confirmed with SDS-PAGE (Figs. S2a and S7a).

#### Preparation of kinesin-streptavidin complexes

Kinesin-streptavidin complexes were prepared by incubating kinesin dimers and streptavidin tetramers in 1xPEM80 buffer (80 mM PIPES, 2 mM MgCl2 and 1 mM EGTA, pH 6.9; supplemented with 5 mM DTT) on ice for 30 min^2^. The solution was then flash frozen in liquid nitrogen, stored at -80°C, and thawed immediately before mass photometry measurements.

For data shown in Figs. 2-4, the final concentration of streptavidin tetramer in the mixture was kept constant at 0.6 µM, the final concentration of kinesin dimers was varied between 0.4-2.16 µM.

For data shown in Fig. S9, the molar ratio of kinesin dimers to streptavidin tetramers was kept constant at 2.1:1. The final concentration of streptavidin tetramers in the mixture was varied between 0.3-2.4 µM, and the final concentrations of kinesin dimers was varied between 0.6-4.8 µM.

Concentrations of isolated proteins were estimated via absorption measurements at 280 nm on a NanoDrop 2000C spectrophotometer (Thermo Scientific). The mixing ratio for each preparation of kinesin-streptavidin complexes was determined via quantitative densitometry of proteins stained with Coomassie blue using an Amersham Typhoon imaging system (GE Healthcare Life Sciences) and quantified via ImageJ (https://imagej.net/ij/) (Fig. S2). Results of these two independent methods agreed to within ∼30%. In our experience, densitometry-based measurements are more reliable and less variable.

#### Mass photometry experiments

All mass photometry experiments were performed at room temperature on a OneMP instrument (Refeyn)^5^. Sample wells were assembled by adhering a 6-well silicone gasket onto a clean coverglass (Refeyn MP-CON-21004). For each set of measurements, 10 µl of protein-free buffer (1xPEM80, supplemented with 5 mM DTT) was added to a new sample well, and the interface between the buffer and the coverglass surface of the sample well was brought into focus. 10 µL of sample (40 nM total protein concentration) was then added to the same sample well, resulting in a final 20 nM total protein concentration. Binding of proteins or protein complexes onto the coverglass surface was then imaged within the 3 μm × 10 μm instrument field of view (Fig. 1b) and recorded for 1 min at a frame rate of 1 kHz. Each sample well was used once. Each preparation of kinesin-streptavidin complexes was measured 12-22 times independently. Reproducibility in mass photometry measurements was verified among independent experiments using the same sample (Fig. S3a) and using different preparations of the same kinesin-streptavidin mixture (Fig. S3b).

#### Data analysis

All mass photometry images were processed and analyzed using DiscoverMP (Refeyn). Interference intensities of protein complexes were converted to masses through a calibration with known mass standards^5-7^ (Fig. S1). The returned molecular masses were presented as mass distribution histograms with 6 kDa bin width. A lower-bound molecular mass of ∼50 kDa was used for each mass distribution to account for the detection lower-bound of the OneMP instrument used in this study^5, 7^. To account for skewness that is common in mass photometry data^6, 8-11^, mass distributions were fitted to a bi-Gaussian mixture model, 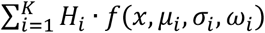, where *K*is the number of major species in the fit, *μ*_*i*_ is the molecular mass of the *i*^th^ mass species, *H*_*i*_ is the peak height of the *i*^th^ mass species, and 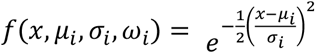 for *x* < *μ*_*i*_, and 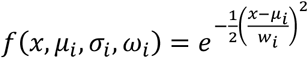 for *x* > *μ*_*i*_ .

The molecular masses of the major species in Fig. 3 were determined via best-fits of mass distributions to the bi-Gaussian mixture model described above. Only mass species with a pronounced peak profile (>40 in peak height, or >400 total counts) were included in the fit (red lines, Fig. 2) to ensure fitting accuracy.

The abundances of the three major complex species (iii-v, Fig. 3) in each mixture were determined via best-fits of mass distributions to the bi-Gaussian mixture model described above. All three major complex species were included in each fit (for example, Fig. S6a). For complex species without a pronounced peak profile, we employed our measurements of molecular masses in Fig. 3 to constrain their peak positions in the fit. The abundance of each major complex species was determined as 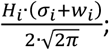 the associated uncertainty was determined via error propagation of fitting uncertainties.

The abundance of the higher-order complexes was determined as the cumulative counts of masses ≥450 kDa (Fig. S4); the associated uncertainty was determined as the counting noise.

Relative abundances of different species of kinesin-streptavidin complexes were determined as the abundance of individual species, normalized by their total abundance. Uncertainties of relative abundances were determined via error propagation. Calculations of relative abundance excluded the isolated proteins. The resulting relative-abundance calculations agreed well between experiments using different preparations of the same kinesin-streptavidin mixture (Fig. S6).

#### Binding model

Monte Carlo simulations were employed to model the distribution of kinesin-streptavidin complexes. Each streptavidin tetramer was modelled as containing four identical sites. Each kinesin dimer was modelled as having up to two identical biotins. Kinesin dimers without a biotin were assumed to not bind streptavidin. A kinesin dimer with one biotin was modelled as a single biotin and could bind a single streptavidin site. A kinesin dimer with two biotins was modelled as two identical biotins coupled together: once the first biotin bound the streptavidin, the second biotin would also bind, provided that there was an open site on the streptavidin. A kinesin dimer with two biotins was assumed to be twice as likely to encounter a streptavidin molecule than a single biotin.

Under these assumptions and denoting *b* as the kinesin-biotinylation efficiency, the fraction of kinesin dimers with two biotins was *b*^2^, the fraction of kinesin dimers with one biotin was 2*b*(1 − *b*), the probability that streptavidin encountering a kinesin dimer with two biotins was *b*, and the probability of streptavidin encountering a kinesin dimer with one biotin was 1 − *b*. At each encounter, the probability that the first (or only) biotin on a kinesin dimer bound a streptavidin site was approximated as the molar ratio of biotin to streptavidin monomers present in the mixture, 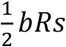, where *b* is the kinesin-biotinylation efficiency denoted above, *R* is the molar mixing ratio determined in experiments, and *s* is an overall scaling factor for the experimentally determined mixing ratios.

Each simulation returned the stoichiometry of a single complex. The simulation was repeated 100000 times to determine the distribution of kinesin-streptavidin complexes for each value of the kinesin-biotinylation efficiency (*b*), the mixing ratio (*R*), and the overall mixing-ratio scaling factor (*s*). Best-fit values of the *b* and *s* parameters were determined by globally minimizing the reduced χ^2^ between the model and the experimental data over all mixing ratios (Fig. S8).

## Supplementary Figures

**Fig. S1.**
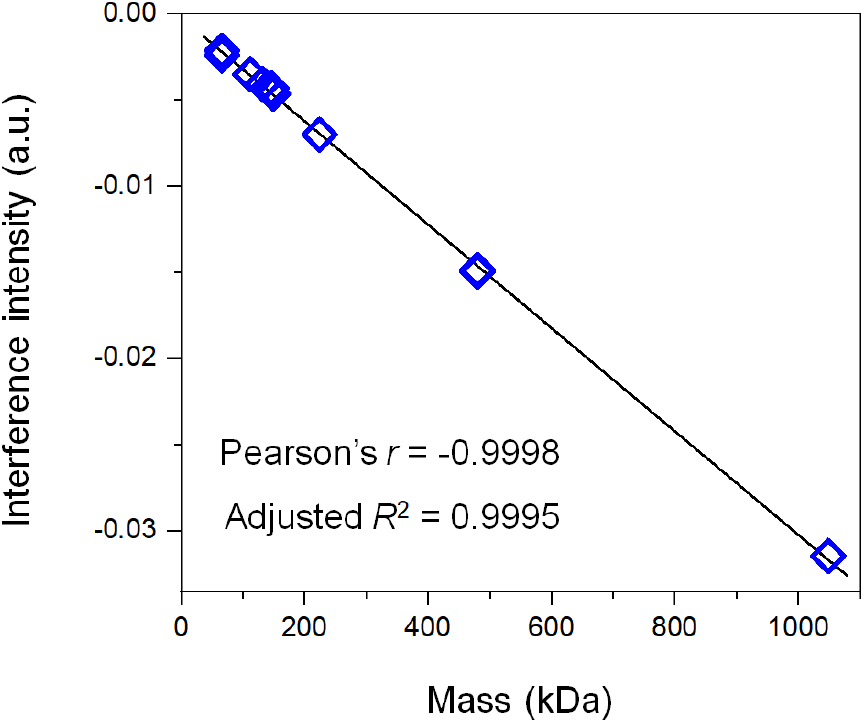
Calibration of molecular mass versus interference intensity for the OneMP instrument (Refeyn, UK)^5-7^. Black line, best linear fit. Pearson’s *r* = -0.9998 and adjusted *R*^2^ = 0.9995.

**Fig. S2.**
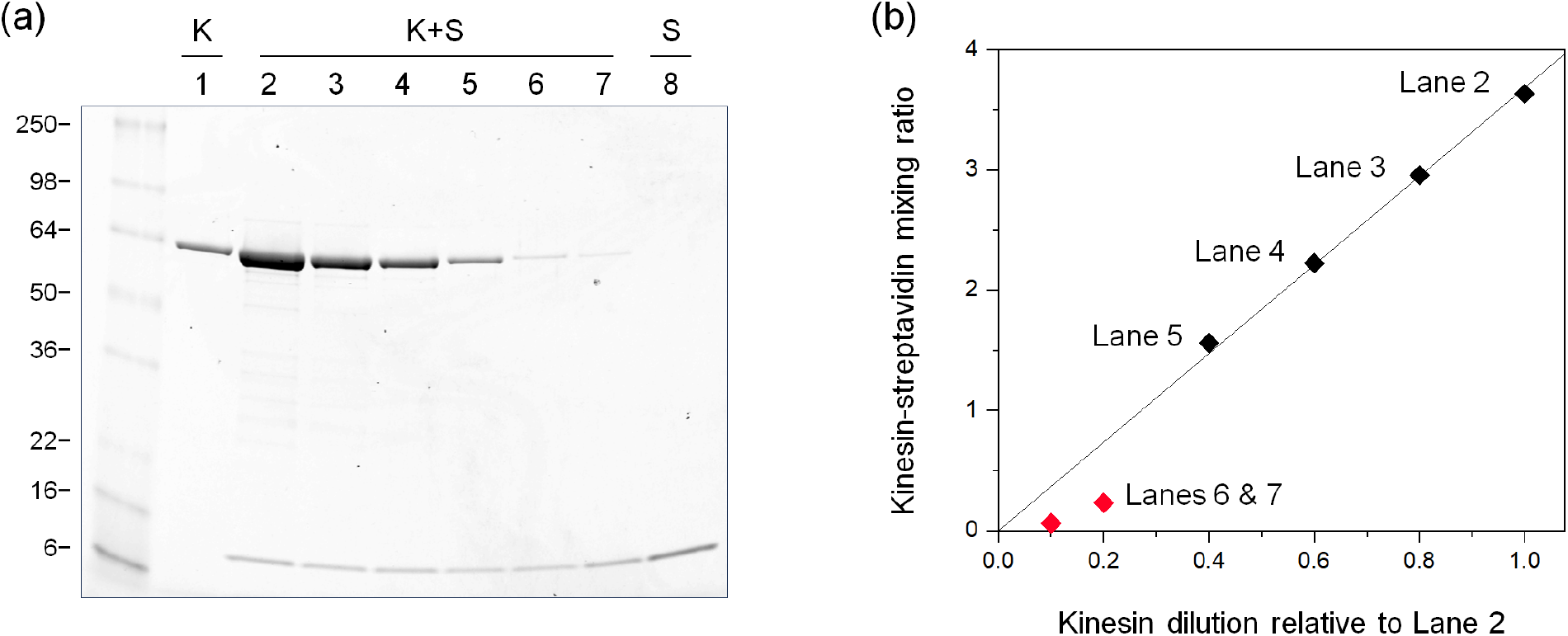
Kinesin-streptavidin mixing ratio determined via quantitative densitometry of proteins stained with Coomassie blue. (a) Denaturing SDS-PAGE gels of kinesin (K), streptavidin (S), and their mixtures (K+S). Positions of molecular weight standards (in kDa) are indicated. (b) The mixing ratio of kinesin dimer to streptavidin tetramer as a function of kinesin dilution relative to Lane 2. Black line, best linear fit with zero intercept. Pearson’s *r* = 0.9998 and adjusted *R*^2^ = 0.9995. For complexes in Lanes 6 & 7, the intensities of kinesin bands in the gel (panel a) were below the linear range of densitometry measurements. These densitometry results were therefore excluded from the linear fit (red points, panel b), and the mixing ratios were estimated via the best-fit result (0.8 and 0.4 kinesin dimer per streptavidin tetramer, respectively).

**Fig. S3.**
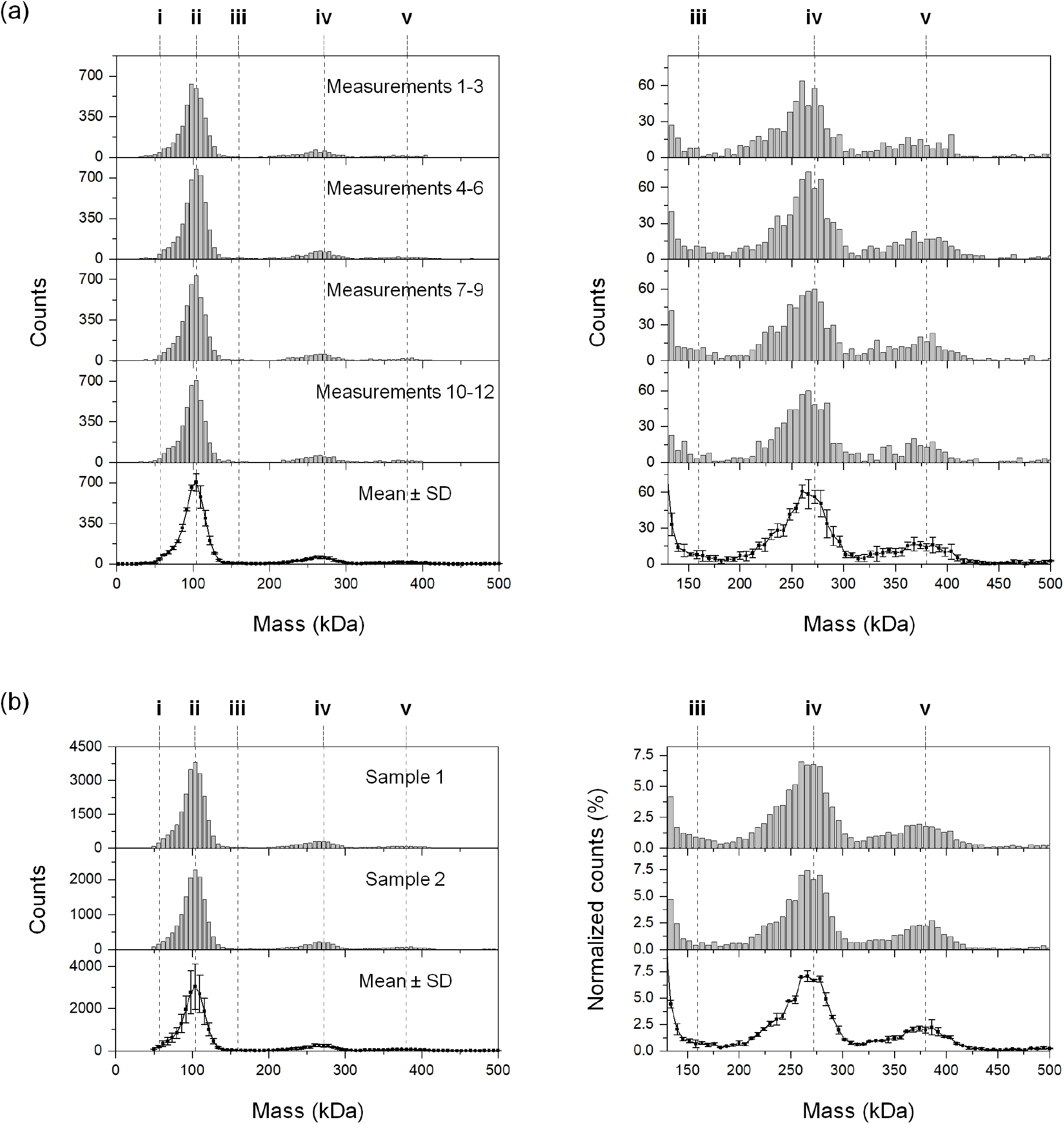
Reproducibility of mass photometry measurements among independent experiments using the same sample preparation (a) and using different preparations of the same kinesin-streptavidin mixture (b). The concentration of streptavidin was kept constant at 0.6 µM. The mixing ratio was kept constant at 3.0 kinesin dimers per streptavidin tetramer. Dashed lines indicate mass species identified in the current study. Expanded view of mass species iii-v are shown on the right. (a) Measurements using the same preparation of kinesin-streptavidin mixture (Lane 3, Fig. S2). Top four panels, mass distributions using data pooled from three independent measurements; measurements were pooled in triplets to increase counting statistics. Bottom panel, mean (± standard deviation) of the mass distributions in the top four panels. (b) Measurements using two different sample preparations of the same kinesin-streptavidin mixture. Sample 1 corresponds to data pooled from the twelve independent measurements shown in (a). Sample 2 corresponds to data pooled from fifteen independent measurements of a second preparation using the same mixing ratio. Mass distributions are shown in raw counts (left) and in counts normalized by the total counts of kinesin-streptavidin complexes (right). Error bars indicate standard deviation.

**Fig. S4.**
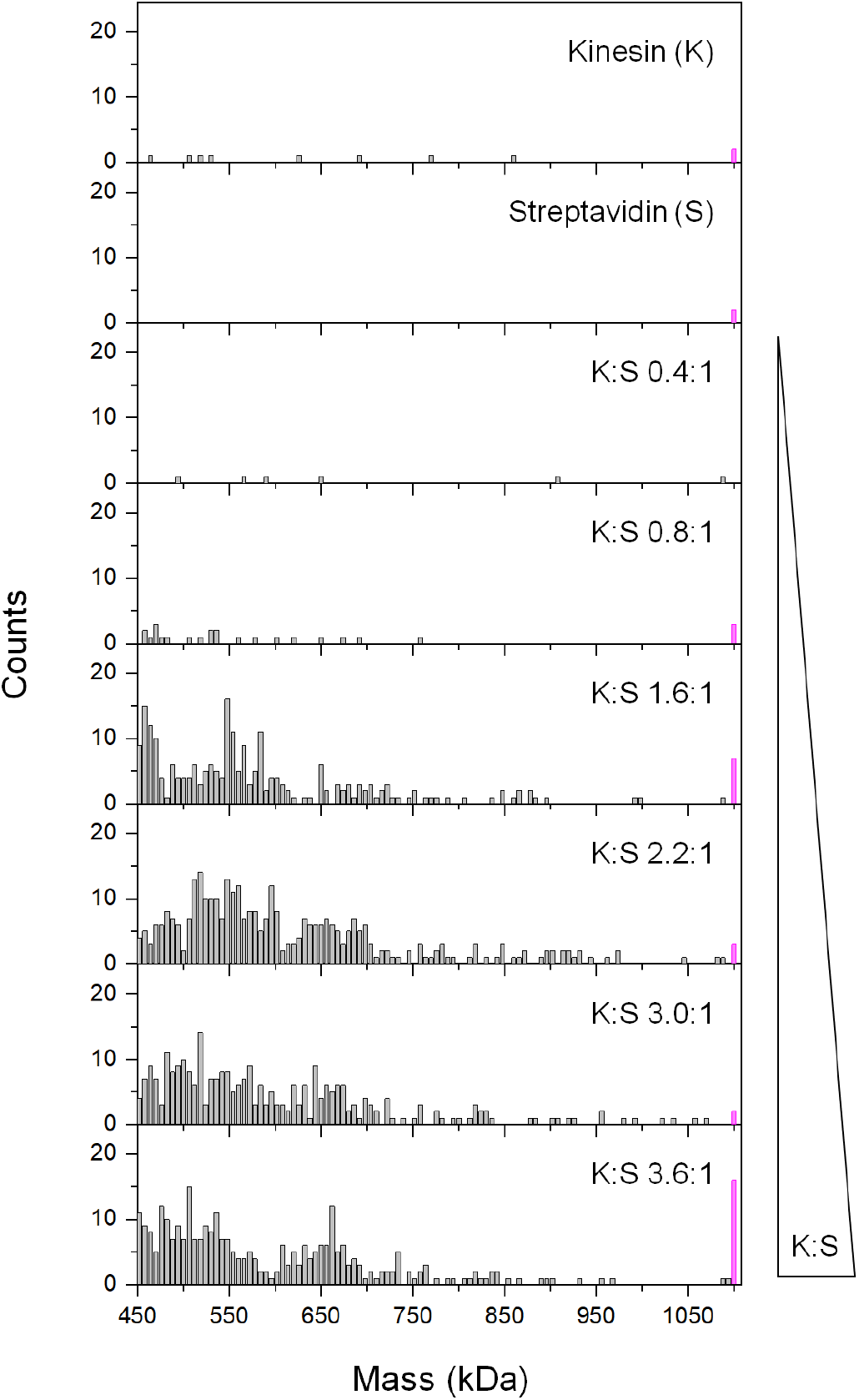
Expanded view of mass distributions from Fig. 2, showing masses larger than 450 kDa for solutions containing kinesin (K), streptavidin (S), and their mixtures (K:S). K:S indicates the molar mixing ratio of kinesin dimers to streptavidin tetramers in each incubation. The concentration of streptavidin tetramers in each mixture was kept constant at 0.6 µM, and the concentration of kinesin dimers was varied. Magenta bars indicate cumulative counts of measurements with masses exceeding 1100 kDa.

**Fig. S5.**
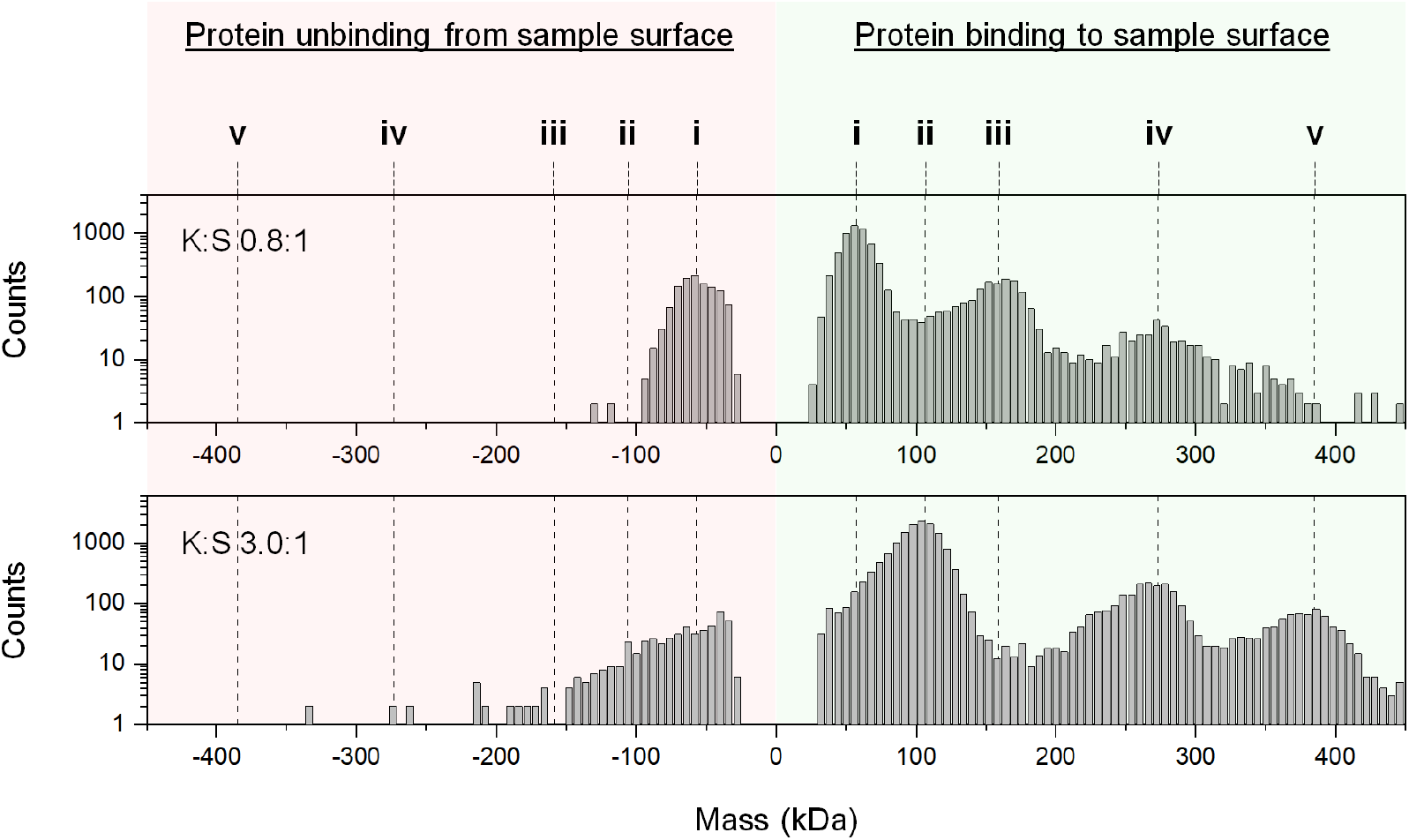
Example mass photometry measurements indicating substantial unbinding of isolated proteins (i-ii), but not kinesin-streptavidin complexes (iii-v), from the sample surface. Negative mass readings (pink region) indicate unbinding events. Positive mass readings (blue region) indicate binding events. The unbound protein can be counted multiple times, through repeated rebinding and unbinding events, artifactually increasing the molecular counts. We therefore excluded the isolated proteins from calculations of the relative abundance of different complex species.

**Fig. S6.**
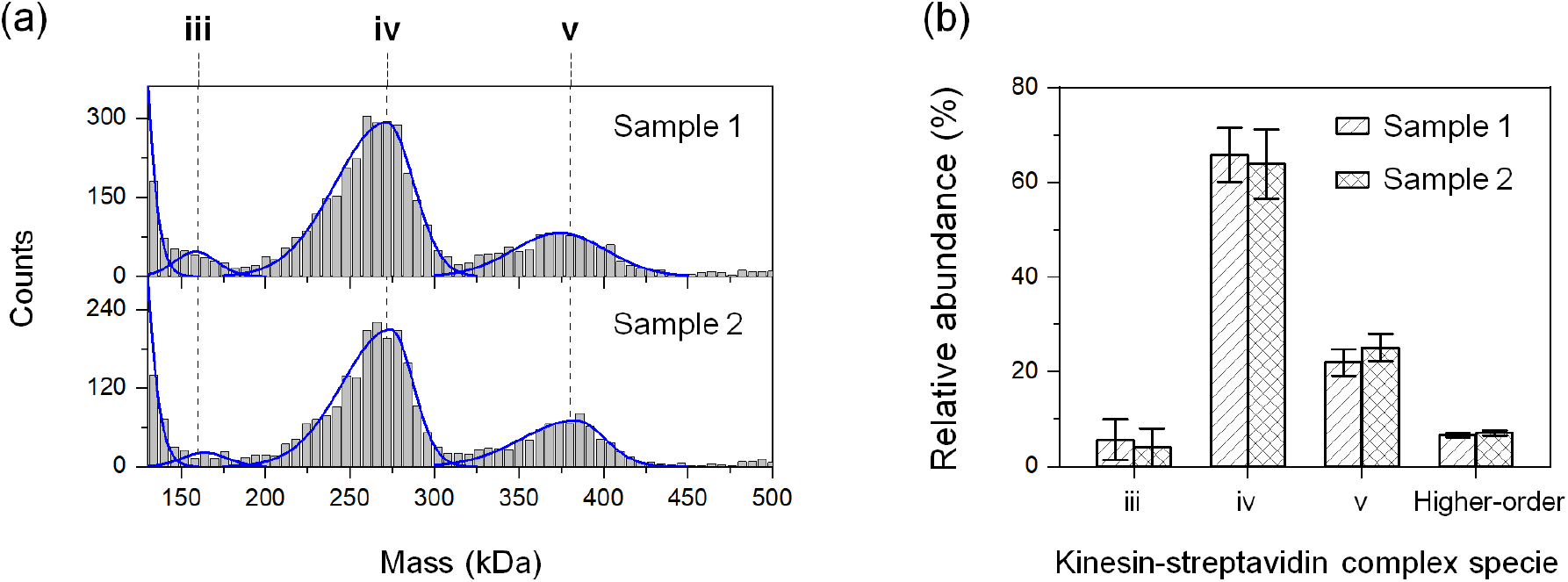
Reproducibility in relative-abundance calculations between experiments using two independent preparations of the same kinesin-streptavidin mixture. The concentration of streptavidin was kept constant at 0.6 µM. The mixing ratio was kept constant at 3.0 kinesin dimers per streptavidin tetramer. (a) Distributions of molecular masses. Blue lines indicate the best-fitted bi-Gaussian peaks for individual mass species. Mass distribution of sample 1 is shown in Fig. 2 (K:S 3.0:1). (b) Relative abundance of complex species determined for the two samples. Calculations of relative abundance excluded the isolated proteins. For complex species iii-v, relative abundances were calculated based on best-fitting results in panel a (blue lines); error bars indicate the associated fitting uncertainties. For higher-order complexes, relative abundances were estimated as the cumulative counts of masses ≥450 kDa; error bars indicate the associated counting noise.

**Fig. S7.**
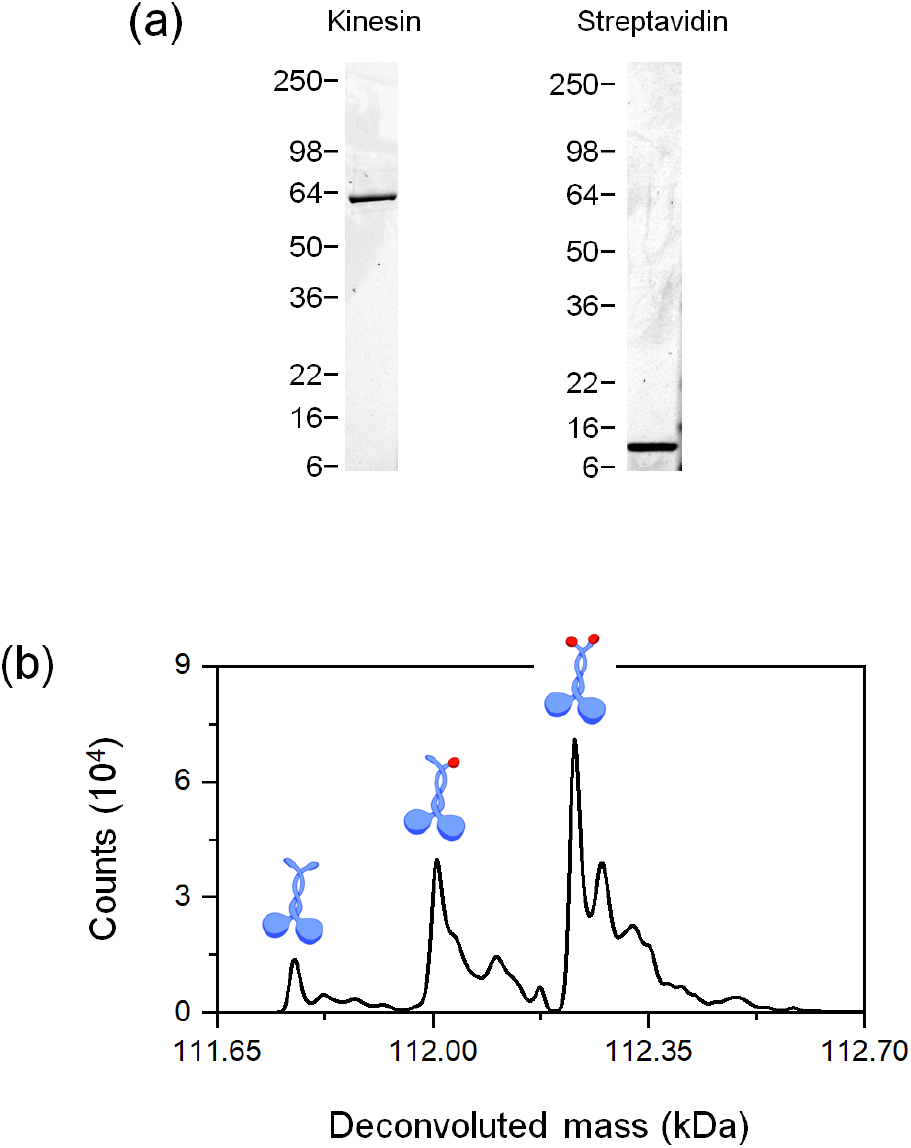
(a) Representative denaturing SDS-PAGE of kinesin and streptavidin proteins used in this study. (b) Liquid chromatography-mass spectrometry (LC-MS) spectra of kinesin. Cartoons illustrate dimeric kinesins (blue) containing 0, 1, and 2 biotins (red). Relative abundances of kinesins with 0 biotin (8%), 1 biotin (30%), and 2 biotins (62%) correspond to ∼77% efficiency of biotinylation of the kinesin monomer.

**Fig. S8.**
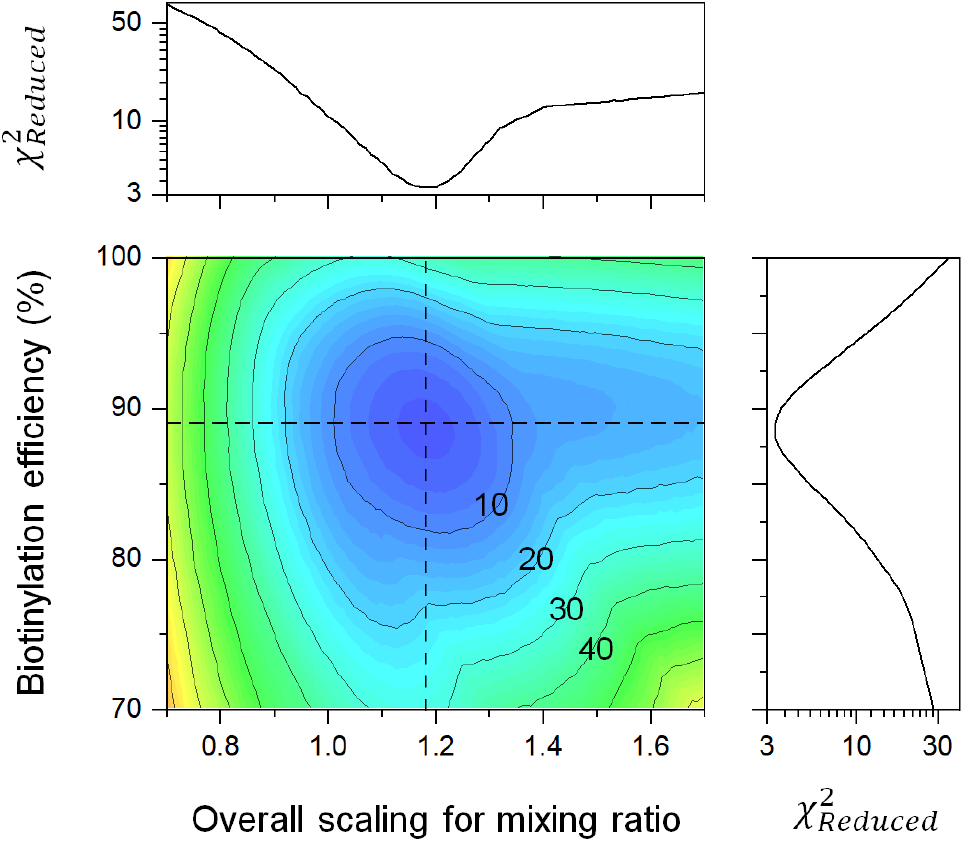
Contour plot of reduced χ^2^ as a function of the kinesin-biotinylation efficiency and the overall scaling for the kinesin-streptavidin mixing ratio in the binding model. Reduced χ^2^ contours of 10, 20, 30, 40 are as indicated. Dashed lines, parameter values that minimize the reduced χ^2^ between the model and the experiments, corresponding to an 88% efficiency in kinesin biotinylation and an overall scaling factor of 1.19 for the kinesin-streptavidin mixing ratio.

**Fig. S9.**
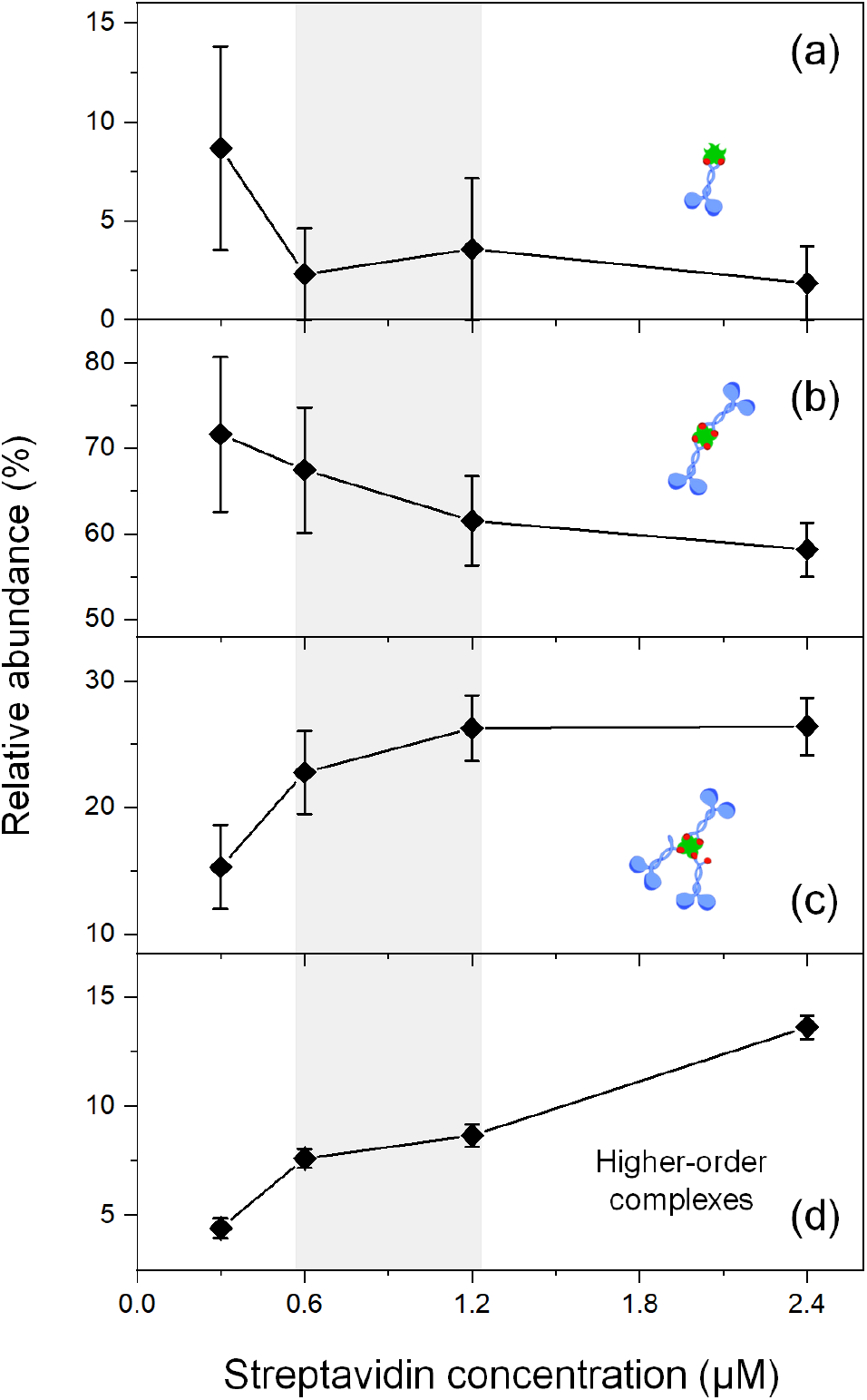
Relative abundance of kinesin-streptavidin complex species as function of the concentration of streptavidin tetramers in the mixture. The mixing ratio was kept constant at 2.1 kinesin dimers per streptavidin tetramer. Calculations of relative abundance excluded the isolated proteins. (a-c) Complexes with 1:1, 2:1, and 3:1 stoichiometry. Relative abundances were determined via best-fits of mass distributions to a bi-Gaussian mixture model; error bars indicate the associated fitting uncertainties. (d) Higher-order complexes. Relative abundances were estimated as the cumulative counts of masses ≥450 kDa in the mass distribution; error bars indicate the associated counting noise. Grey region, the relative abundance of complex species remained largely unchanged for streptavidin concentrations ranging between ∼0.6-1.2 µM.

**Fig. S10.**
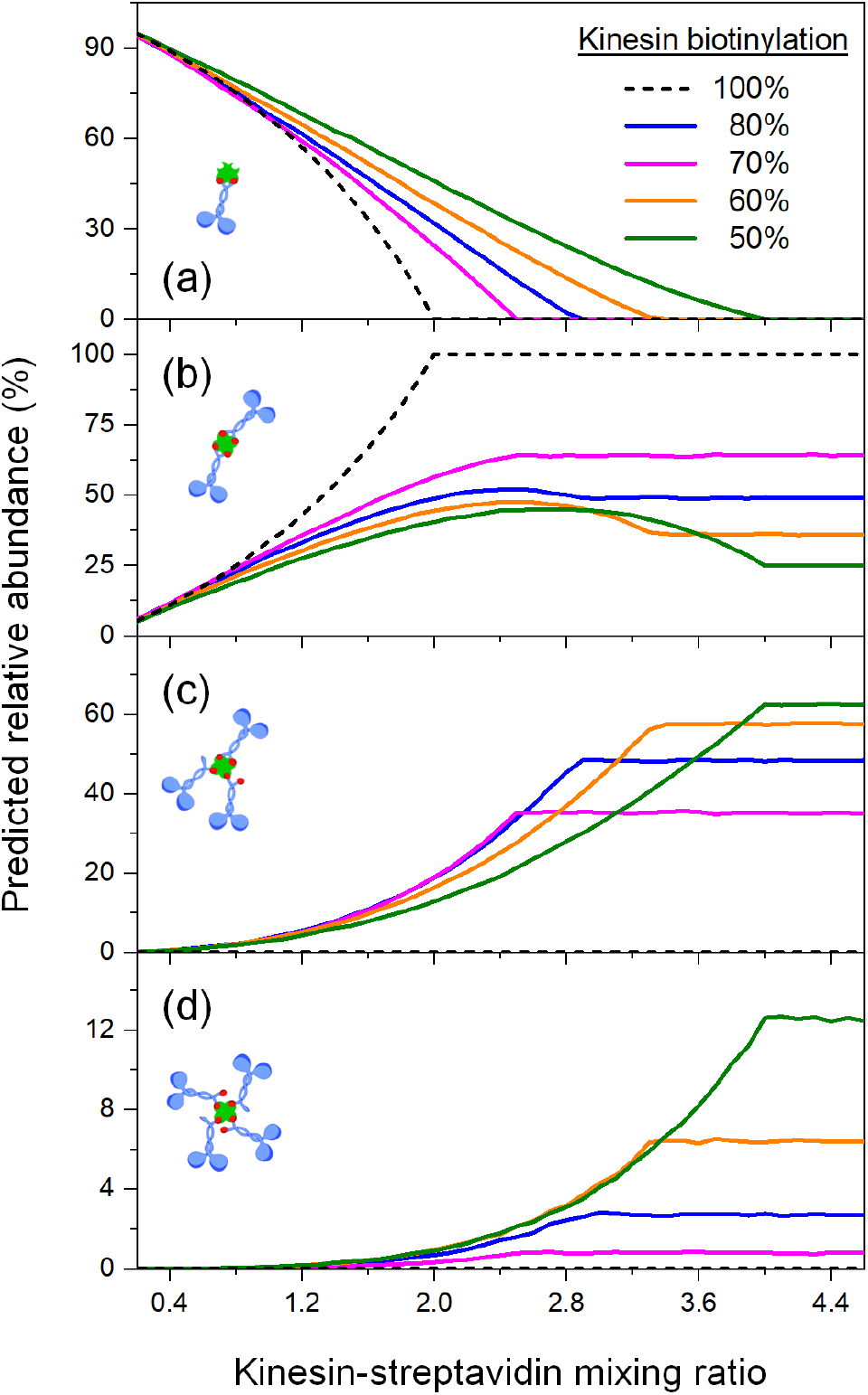
Predicted relative abundance of kinesin-streptavidin complex species as function of the kinesin-streptavidin mixing ratio for five different kinesin-biotinylation efficiencies. Note that our simple binding model makes explicit the relative abundances of complexes with well-defined stoichiometries (a-d), but does not consider higher-order complexes. Calculations of relative abundance excluded the isolated proteins.

